# Deep evolutionary origin of limb and fin regeneration

**DOI:** 10.1101/503011

**Authors:** Sylvain Darnet, Aline C. Dragalzew, Danielson B. Amaral, Andrew W. Thompson, Amanda N. Cass, Jamily Lorena, Josane de Freitas Sousa, Carinne M. Costa, Marcos P. Sousa, Nadia B. Fröbisch, Patricia N. Schneider, Marcus C. Davis, Ingo Braasch, Igor Schneider

## Abstract

Salamanders and lungfishes are the only sarcopterygians (lobe-finned vertebrates) capable of complete limb and paired fin regeneration, respectively. Among actinopterygians (ray-finned fishes), regeneration after amputation at the fin endoskeleton has only been demonstrated in Polypterid fishes (Cladistia). Whether complete appendage regeneration in sarcopterygians and actinopterygians evolved independently or has a common origin remains unknown. Here we combine fin regeneration assays and comparative RNA-seq analysis to provide support for a common origin of a paired appendage regeneration in osteichthyes (bony vertebrates). We show that, in addition to Polypterids, regeneration after fin endoskeleton amputation occurs in extant representatives of all major actinopterygian clades: the American paddlefish, (Chondrostei), the spotted gar (Holostei), as well as in two cichlid species, the white convict and the oscar (Teleostei). Our comparative RNA-seq analysis of regenerating blastemas of axolotl and *Polypterus* reveals the activation of common genetic pathways and expression profiles, consistent with a pan-osteichthyes genetic program of appendage regeneration. Collectively, our findings support a deep evolutionary origin of paired appendage regeneration in osteichthyes and provide an evolutionary framework for studies on the genetic basis of appendage regeneration.

---

A typical osteichthyan paired fin (i.e. pectoral and pelvic fin) is composed of an array of proximal fin radials (the endoskeleton), followed distally by the fin rays (the dermal skeleton). The tetrapod limb evolved from paired fins in Devonian sarcopterygians. During this transition, the fin ray dermal skeleton was lost and the elaborate limb endoskeleton emerged, consisting of a proximal segment, the stylopod (humerus and femur), an intermediate segment, the zeugopod (radius/ulna, tibia/fibula), and a distal segment, the autopod (manus and pes)^1^. Therefore, whereas fin rays have no direct homologous counterpart in tetrapod limbs, the limb endoskeleton and the endoskeletal elements of fish paired fins share deep homology^2,3^ (Fig. 1).

**Fig. 1.**
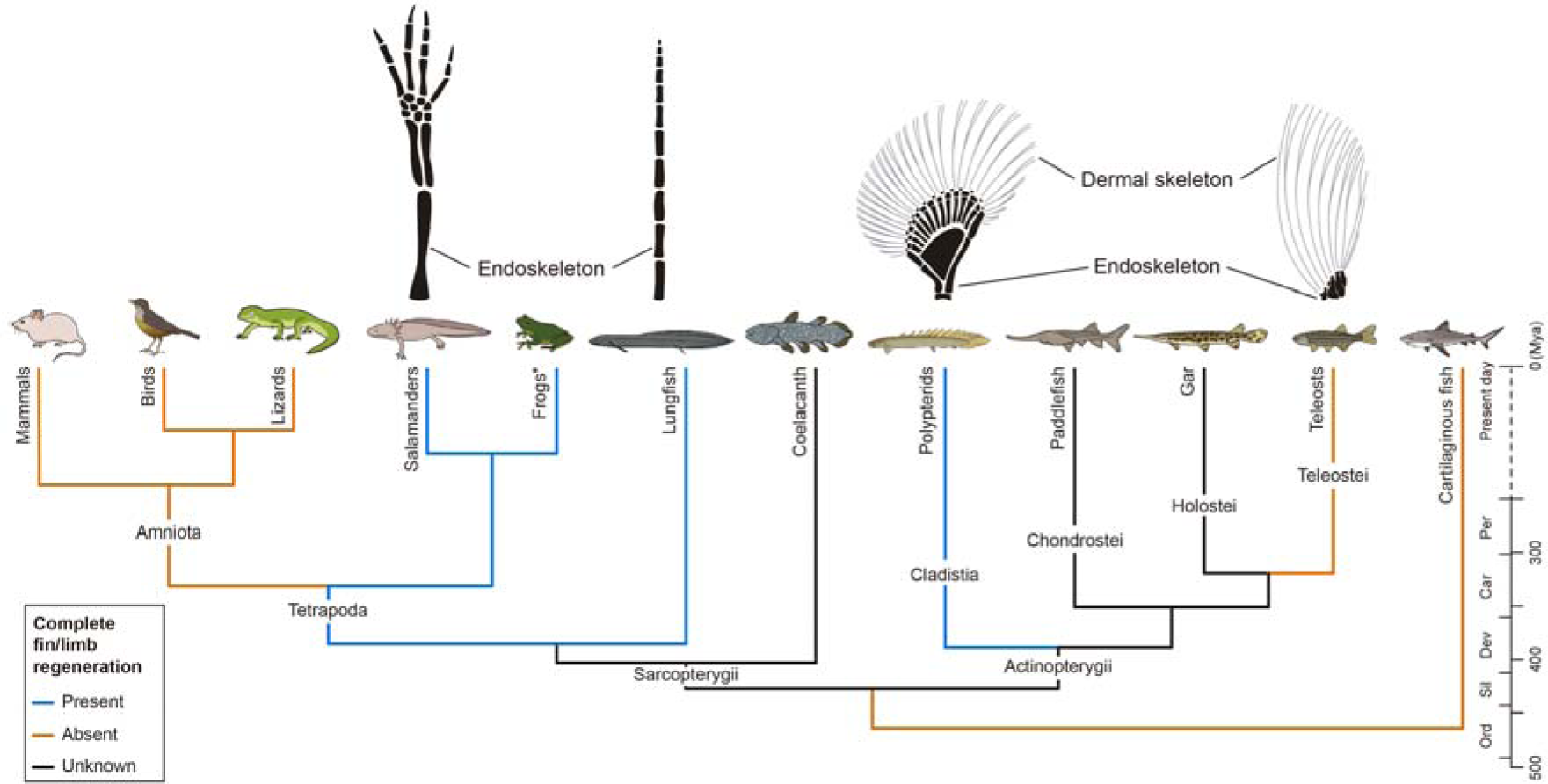
Phylogenetic distribution of complete appendage regeneration among vertebrates. Top: schematic representation of endoskeleton and dermal skeleton of vertebrate appendages. Bottom: a time-calibrated vertebrate phylogeny depicts the state of current knowledge at the time this study was undertaken. Lineages containing species capable of complete fin or limb regeneration are denoted in blue, those incapable denoted in orange and those where no information exists in black.

Among sarcopterygians, the capacity to regenerate the limbs and fins after amputations severing the endoskeleton has been reported only in three groups: frogs^4^, salamanders^5^ and lungfishes^6^. Although adult frogs cannot regenerate limbs, this capacity is exhibited by tadpoles prior to metamorphosis^7^. Recent fossil evidence showed that limb regeneration occurred in basal amphibians prior to the emergence of stem salamanders, caecilians and frogs, hence this capacity is likely an ancient, plesiomorphic feature of tetrapods^8,9^. Recently, a transcriptome analysis revealed strong similarities between the transcriptional profiles deployed in lungfish fin and salamander limb blastemas^6^. Altogether, current data support the hypothesis that tetrapods inherited a limb regeneration program from sarcopterygian fish ancestors^10^.

Among actinopterygians, teleosts such as zebrafish have been broadly used for fin regeneration studies. However, their regenerative abilities are thought to be limited to the dermal fin ray skeleton^11,12^. Thus far, only two actinopterygian species, both from the Polypteridae family, have been found to fully regenerate paired fins including the endoskeleton: the Senegal bichir *Polypterus senegalus*^13,14^ and the ropefish *Erpetoichthys calabaricus*^14^. Currently, our understanding of the evolution of appendage regeneration is hindered by limited knowledge of the regeneration capabilities across fish species. To address this, we assessed fin regeneration capacity in key taxa representing all extant major actinopterygian clades and examined gene expression profiles of limb and fin regenerating blastemas via RNA-seq.

Here we provide evidence of regeneration after amputation at the fin endoskeleton in the American paddlefish, (Chondrostei), the spotted gar (Holostei), and in two cichlid species, (Teleostei), which together with Polypterids (Cladistia) constitute living representatives of all major actinopterygian lineages. Further, we show that regenerating blastemas of axolotl and *Polypterus* activate common genetic pathways and expression profiles, including shared expression of regeneration-specific genes. Altogether, these findings suggest that regeneration of paired fins and limbs in modern vertebrates have a common deep evolutionary origin.

## Results

The American paddlefish (*Polyodon spathula*) is a descendant of the early-diverging actinopterygian clade Chondrostei, (Acipenseriformes) (Fig. 1). Paired fins of paddlefish are supported proximally by an elaborate endoskeleton. We chose to assess regeneration of pelvic fins, which have an endoskeleton compartment more readily accessible to amputations (Fig. 2a). We performed a total of 8 pelvic fin amputations of juvenile fish and assessed for regenerative growth 4 weeks later (Fig. 2b). We found that at 28 days post-amputation (dpa), 6 of 8 fish showed chondrogenic outgrowth and repatterning distal to the amputation plane (Fig. 2c). All specimens displayed heteromorphic regeneration, where the regenerated endoskeleton and dermal skeleton morphology differed from the original, with significant bifurcations of radials occurring in register with the amputation plane, as well as novel condensations and bars of cartilage (Fig 2c, Supplementary Fig. 1a-d). In 4 of 6 fish with regenerative outgrowths there was significant regrowth of the dermal fin-fold including the formation of lepidotrichia. In sum, these results showed that juvenile paddlefish are capable of fin regeneration after amputation at the fin endoskeleton.

**Fig. 2.**
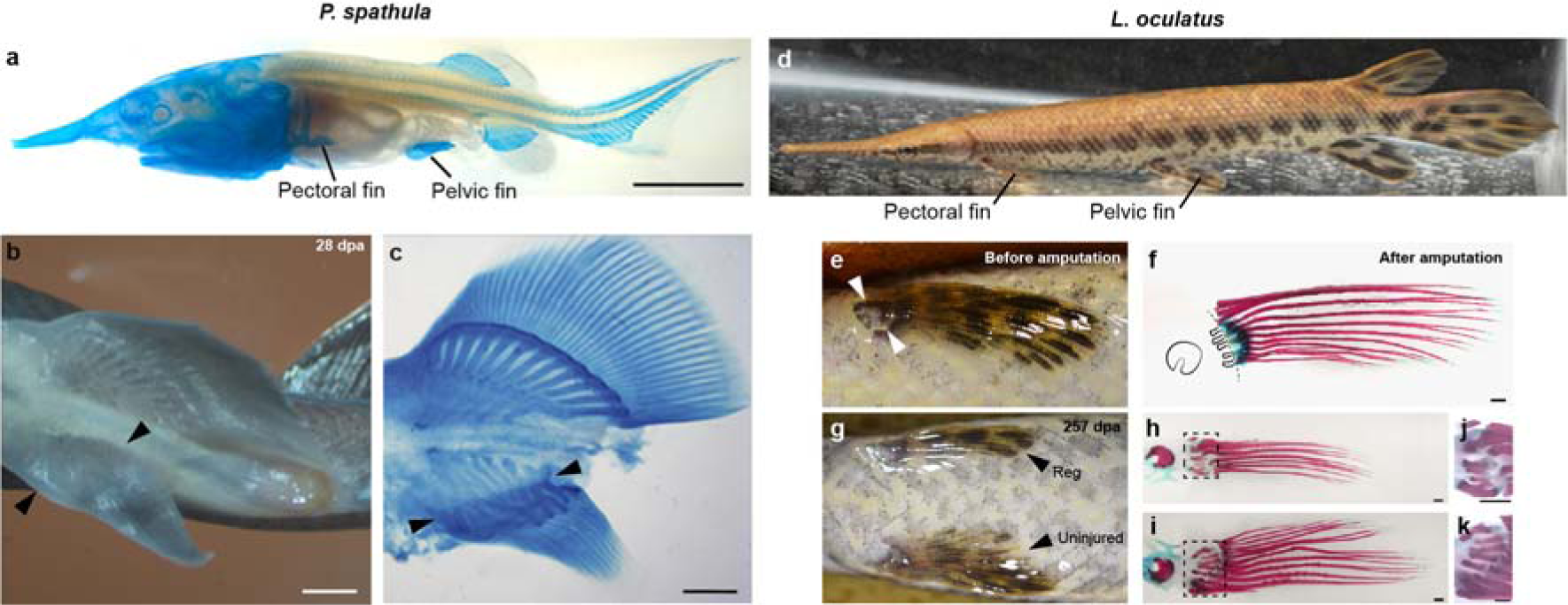
Evidence of regeneration after fin endoskeletal amputation in non-teleost actinopterygians. **a**, Cleared and stained juvenile paddlefish at 75 days post-fertilization (dpf); pectoral and pelvic fins denoted. **b**, Ventral view of a specimen with regenerated right pelvic fin at 28 dpa; arrowheads denote amputation site. **c**, Cleared and stained specimen showing endoskeletal regeneration distal to the amputation site. **d**, Spotted gar with pectoral and pelvic fins denoted. **e**, Left pelvic fin before amputation. **f**, Skeletal staining of fin removed by amputation; dotted line shows amputation site across the endoskeleton. **g**, Ventral view showing regenerated left fin at 257 dpa and uncut right fin. **h**, Skeletal staining of regenerated fin. **j**, Close-up view of regenerated endoskeleton. **i**, Skeletal staining of uncut right fin. **j**, Close-up view of uncut right fin endoskeleton. Scale bars of 5 mm (**a**) and 1 mm (**b** and **c**, **f**, **h-k**). In all panels anterior is to the left.

Gars are members of the Lepisosteiformes (Holostei) and together with Polypterids and the chondrostean paddlefish, constitute living representatives of the three principal non-teleost clades of living actinopterygians^15^ (Fig. 1). We performed pectoral fin amputations across the endoskeleton (Fig. 2d-f) on 15 individuals and followed regenerative outgrowth for over 8 months. A total of 11 of 15 fish displayed various degrees of regeneration, from partial to near complete regrowth, and the regenerated fin radials and rays that were mostly shorter and misshapen (Fig. 2g-k, Supplementary Fig. 1e-h). Therefore, as seen in paddlefish, regeneration was mostly heteromorphic. Nevertheless, collectively, our results on gar and paddlefish and previous reports in Polypterids suggest that the capacity for regeneration after fin endoskeleton amputation is a common feature of living non-teleost actinopterygian.

Given the observations above, the question of whether complete fin regeneration could extend to the teleosts was reexamined. To this end, we selected two cichlid species, the white convict *(Amatitlania nigrofasciata)* and the oscar *(Astronotus ocellatus)*, in which the pectoral fin endoskeleton compartment was sufficiently large and accessible for amputations (Fig. 3a, f). As seen in gar, regeneration after amputation at the fin endoskeleton progressed slowly, and was followed for several months. At 160 dpa, fin regeneration was observed in 6 of 8 white convicts (Fig. 3b, Supplementary Fig. 2a, e). Likewise, 3 of 4 oscars showed fin regeneration at 90 dpa (Fig. 3g, Supplementary Fig. 2i, m). In both species, the extent of regeneration varied, and regenerated fins differed from the original morphology. Skeletal staining of the amputated fins confirmed that amputation plane crossed the fin endoskeleton, removing the distal ends of the radials (Fig. 3 c, h and Supplementary Fig. 2b, f, j, n). In white cichlids, regenerated fin radials displayed discrete distal outgrowth and some radials partially recovered the original morphology (Fig. 3d, e and Supplementary Fig. 2c, g). Fin radial distal outgrowth was occasionally associated with hypertrophy (Fig. 3e and Supplementary Fig. 2d, h). The regenerated dermal skeleton was characterized by fin rays that were shorter and reduced in number (Fig. 3d and Supplementary Fig. 2c, g). In oscars, regenerated fin radials also showed distal outgrowth and hypertrophy (Fig. 3i, j and Supplementary Fig. 2k, l, o, p), and fin rays were shorter and reduced in number. Altogether, these results demonstrated that despite being predominantly heteromorphic, regeneration of paired fins following amputation through the endoskeleton is observed in representatives all major lineages of living actinopterygians.

**Fig. 3.**
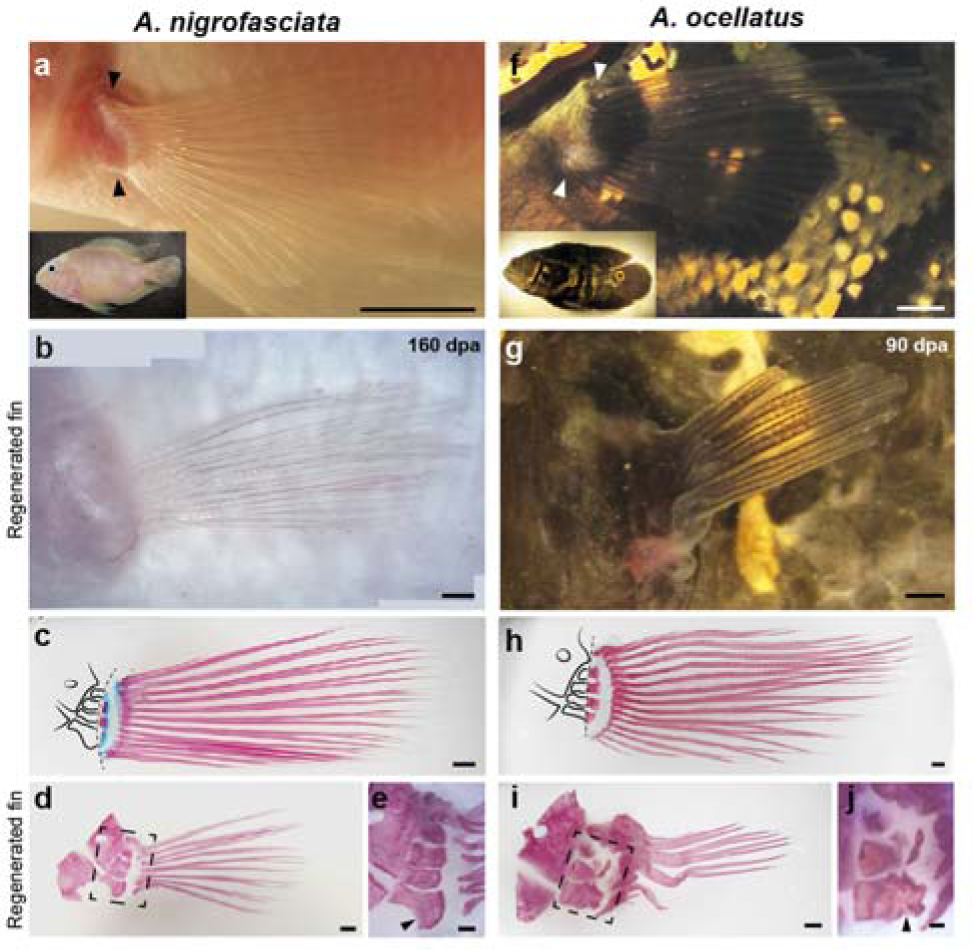
Evidence of regeneration after fin endoskeletal amputation in cichlids. **a**, Side view, white convict specimen (inset) and its left pectoral fin; arrowheads denote amputation site. **b**, Regenerated fin at 160 dpa. **c**, Skeletal staining of fin removed by amputation; dotted line denotes amputation site across the endoskeleton. **d**, Skeletal staining of regenerated fin at 160 dpa. **e**, Close-up view of regenerated endoskeleton. **f**, Side view, oscar specimen (inset) and its left pectoral fin; arrowheads denote amputation site. **g**, Regenerated fin at 90 dpa. **h**, Skeletal staining of fin removed by amputation; dotted line shows amputation site across the endoskeleton. **i**, Skeletal staining of regenerated fin at 90 dpa. **j**, Close-up view of regenerated endoskeleton. Scale bars of 5mm (**a** and **f**) and 1 mm (**b**-**e** and **g**-**j**).

Our findings are consistent with the hypothesis that complete appendage regeneration evolved in osteichthyes before the divergence of the actinopterygian and sarcopterygian lineages. We thus hypothesized that actinopterygian fins and salamander limbs may share a common, ancient genetic program for appendage regeneration. To examine this, we generated RNA-seq data from *Polypterus* fin blastema (FB) and non-regenerating fin (NRF), and from axolotl limb blastema (LB) and non-regenerating limb (NRL) (Supplementary Fig. 3). Spearman correlation coefficients among biological replicas were greater than 0.71, corroborating the reproducibility of RNA-seq runs (Supplementary Fig. 3e).

Differential gene expression (DGE) analysis of the axolotl blastema *versus* NRL revealed 562 downregulated and 1443 upregulated genes. Our axolotl DGE data correlates well to publicly available axolotl limb regeneration RNA-seq profiles, with up and downregulated genes showing equivalent TPM values in all RNA-seq replicas^16^ (Supplementary Table 1 and Supplementary Fig. 4a and b). Next, we performed DGE analysis of the *Polypterus* blastema versus NRF and found 379 downregulated and 957 upregulated genes, including genes typically downregulated (*Mybpc2*, *Casq1*, *Myoz1*, *Smpx*, *Tnnt3*) or upregulated (*Mmp11*, *Sall4*, *Msx2*, *Sp9*, *Wnt5a*, *Fgf8* and *Fgf10*) in axolotl blastemas^17-21^ (Fig. 4a and Supplementary Table 2). Quantitative PCR (qPCR) profiles of 12 differentially expressed targets were largely consistent with the *Polypterus* RNA-seq data (Supplementary Fig 4c). Comparison between axolotl and *Polypterus* blastema DGE datasets revealed that 35.31% of the genes upregulated in the *Polypterus* fin blastema possess homologs upregulated in the axolotl limb blastema (Fig. 4b). Analysis of enriched gene ontology (GO) categories showed that *Polypterus* blastema are enriched for several GO terms associated with axolotl limb blastema, including appendage morphogenesis, extracellular matrix organization and chromatin remodeling (Fig. 4b and Supplementary Table 3). Further, we found that 179 of 265 (67.54%) of the enriched GO categories in the *Polypterus* blastema were also enriched in the axolotl blastema transcriptome (Fig. 4b).

**Fig. 4.**
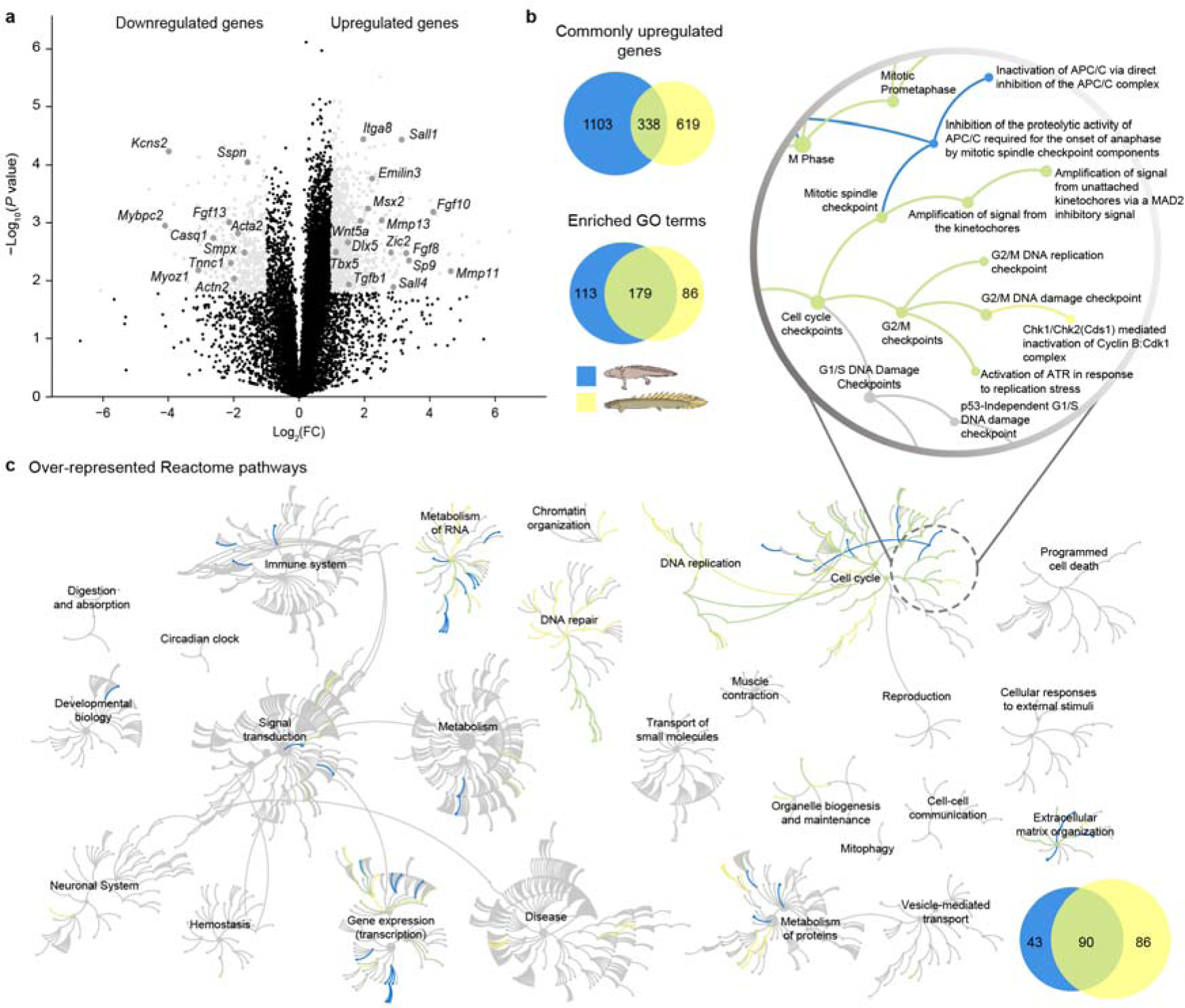
Comparative transcriptome analysis of *Polypterus* fin and axolotl limb blastema. **a**, Volcano plot showing differentially expressed genes in *Polypterus*, between NRF tissue and 9 dpa FB (*P* value<0.05, FC>2). *Polypterus* orthologs of genes commonly up or downregulated in axolotl blastema are pointed out. **b**, Area-proportional Venn diagrams showing upregulated genes (*P* value<0.05, FC>2) and enriched GO categories (*P* value<0.05) in axolotl and *Polypterus* DGE datasets. **c**, A graphical overview of Reactome pathway analysis for axolotl and *Polypterus*, each of the central circles is a top-level pathway, and each step away from the center represents a lower level in the pathway hierarchy (see top right for zoom of a section of the top-level pathway Cell cyle). Over-represented pathways (*P* value<0.05) are colored in yellow (*Polypterus*), blue (axolotl) or green (overlayed pathways from both species), light grey represents pathways not significantly over-represented, an inset (bottom right) shows the Area-proportional Venn diagram of the enriched pathways in both species.

Next, we performed a pathway over-representation analysis on the blastema upregulated genes in *Polypterus* and axolotl (Supplementary Table 4). A graphical representation of this data shows each top-level pathway as a central circle, connected to other circles representing the next level lower in the pathway hierarchy (Fig 4c, see zoom in on a section of the Cell Cycle pathway). Our analysis revealed that 88.1% (155 of 176) and 88.7% (118 of 133) of enriched pathways in *Polypterus* and axolotl, respectively, fall into 7 of 26 broader categories, namely Extracellular matrix organization, Cell cycle, DNA replication, DNA repair, Metabolism of proteins, Metabolism of RNA and Gene expression (transcription) (Fig. 4c). We found that 90 of 133 (67.7%) over-represented pathways in axolotl limb blastema were shared with the *Polypterus* fin blastema dataset (Fig. 4c), including pathways involved in Collagen formation, Extracellular matrix organization, Regulation of TP53 activity and Cell cycle. Conversely, among downregulated genes, we found that 14 of 28 (50%) shared over-represented pathways between *Polypterus* and axolotl, including pathways involved in Muscle contraction and Metabolism (Supplementary Fig. 5). Collectively, our findings revealed substantial similarities of gene expression, GO enrichment and pathway over-representation profiles between *Polypterus* and axolotl blastemas.

**Fig. 5.**
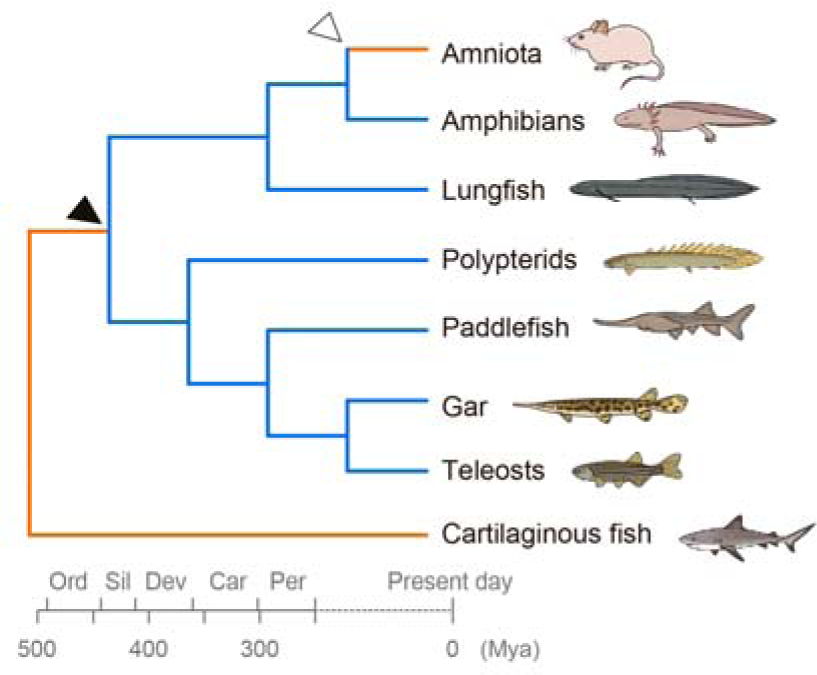
Hypothesis for the evolution of complete limb and fin regeneration in vertebrates. Regeneration-incompetent taxa shown in orange, regeneration-competent taxa in blue; black arrowhead indicates the origin of paired fin regeneration and white arrowhead loss of limb regeneration.

Many molecular networks orchestrating limb development are also redeployed during limb regeneration^22^. In fact, a recent single cell RNA-seq (scRNA-seq) analysis comparing axolotl limb blastemas to developing limb buds has found a high correlation between gene expression profiles of late stage limb blastemas and limb buds^23^. However, the study also revealed that early-stage limb blastemas possess a regeneration-specific gene expression profile distinct from that of developing limb buds and from uninjured limbs. As a result, we sought to determine whether our *Polypterus* blastema DGE dataset contained genes differentially expressed during axolotl limb regeneration relative to limb buds and to uninjured limbs. To this end, we screened the axolotl scRNA-seq dataset for the orthologs of the 957 genes we had identified as upregulated in *Polypterus* fin blastema and found that 862 were present in the axolotl scRNA-seq dataset. Next, we compared the expression levels of these genes in the axolotl scRNA-seq data corresponding to 9 conditions: 5 blastema stages, 3 developmental stages and uninjured limb. We found that 110 of the 862 genes were differentially expressed during at least one stage of regeneration when compared to developing limb buds and to uninjured limbs (Supplementary Fig. 6). This list includes genes found upregulated or previously implicated in appendage regeneration in axolotls and lungfish, such as *Fmod*, *Emilin1*, *Adamts17*, *Mmp11*, *Mmp13*, *Steap1*, *Itga8* and *Tgfb1*^6,21^. This suggests that a genetic program distinctive for axolotl limb regeneration is also found among upregulated genes in the 9 dpa *Polypterus* blastema, consistent with the hypothesis of an ancient genetic program of appendage regeneration shared among osteichthyes.

## Discussion

Heteromorphic regeneration has been previously reported for salamanders^24^, lungfish^6^ and for dermal fin ray regeneration in teleosts^25^. Repeated fin amputations also results in abnormal endoskeleton morphologies in *Polypterus*^14^. In our study, all five species examined showed varying degrees of heteromorphic regeneration, including no regeneration. These findings suggest that a great variability of fin regenerating capacity may exist among actinopterygians. Nevertheless, our results indicate that regenerative outgrowth may be the most common outcome of complete fin amputation.

Further, we showed two teleost species representing lineages that split over 40 mya^26^, in which amputation at the pectoral fin endoskeleton resulted in heteromorphic fin regeneration. The identification of additional teleost species capable of complete fin regeneration may provide a valuable source of novel model species for comparative studies of appendage regeneration.

Finally, the search for genes linked to limb regeneration has been mostly pursued without a well-founded evolutionary context. If limb regeneration has an ancient evolutionary origin, then genes facilitating regeneration in extant species are likely part of an ancient genetic program whereas species without the capacity likely lost this ability over evolutionary time. Therefore, an evolutionarily informed approach based on comparative analysis of regenerating limbs and fins will offer a more powerful method to identify a shared genetic program underlying vertebrate appendage regeneration.

## Methods

### Animal work

*Polypterus* (*Polypterus senegalus*), oscar (*Astronotus ocellatus*) and white convict cichlid (*Amatitlania nigrofasciata*) were maintained in individual tanks in a recirculating freshwater system at 26°C with aeration and experiments and animal care were performed in accordance with animal care guidelines approved by the Animal Care Committee at the Universidade Federal do Para (protocol number 037-2015). Axolotls (*Ambystoma mexicanum*) were obtained from the Center for Regenerative Therapies (Dresden, Germany) and maintained in accordance with the animal care guidelines at the Museum für Naturkunde Berlin (Germany). Paddlefish (*Polyodon spathula*) embryos were obtained from Osage Beach Catfisheries Inc. (Osage Beach, MO, USA), and were raised at 18°C in recirculating large-volume freshwater tanks, in accordance with approved Institutional Animal Care and Use Committee (IACUC) protocols at Kennesaw State University (protocol number 16-001; former institution of M.C.D.), and James Madison University (protocol number A19-02). Spotted gar (*Lepisosteus oculatus*) were obtained as embryos from hormone-induced spawns of wild-caught broodstock from bayous near Thibodaux, LA, USA, raised in 150-300-gallon tanks, in accordance with approved Institutional Animal Care and Use Committee (IACUC) protocols at Michigan State University (protocol number AUF 10/16-179-00).

*Polypterus* were kept in individual tanks with regular water changes and fed once per day. Eighteen fish ranging 5-8 cm were used in the study. Fish were anesthetized in 0.1% MS-222 (Sigma) and pectoral fins were bilaterally amputated across the endoskeleton. A portion of the amputated fins, encompassing the endoskeleton elements, was sampled and labeled as NRF tissue (n=3), and regenerating fin tissue was sampled from the left and right pectoral fins at 1 (n=9, three pools of three individuals each), 5 (n=3) and 9 dpa (n=6). Fish were euthanized in 300mg/L of MS-222 (Sigma). All tissue collected was stored in RNAlater (Sigma) for RNA extraction and subsequent qPCR or RNA-seq experiments.

A total of 6 axolotls ranging from 8-12 cm were used in the study. Animals were kept in individual tanks with regular water changes and fed once per day. For limb regeneration studies, axolotls were anesthetized in 0.1% MS-222 (Sigma) and forelimbs were bilaterally amputated at the level of the upper arm. A portion of the upper arm tissue was sampled and labeled as NRL, and regenerating limb tissue was sampled from the left and right forelimbs at 14 dpa (n=6, three pools of two individuals each). All tissues collected were stored in RNAlater (Sigma) for RNA extraction and subsequent RNA-seq experiment.

Paddlefish were kept in large volume, high flow systems that mimic their natural environment and fed brine shrimp daily. For fin regeneration studies, 8 juvenile fish were anesthetized in 0.1% MS-222 (Sigma) at 48 days post-fertilization (dpf) (a stage when endochondral elements are present in the fin) and pelvic fin amputations were performed. Fish were raised for 28 dpa, to a total length of 8-13 cm, then euthanized with a lethal dose of MS-222 (Sigma), fixed in 4% paraformaldehyde (PFA) in phosphate buffered saline (PBS), and stored in methanol at -20°C until analysis. Specimens were cleared and stained as previously described^27^ and photographed with a Zeiss SteREO Discovery.V12 microscope with MRc5 camera.

Experimental spotted gar were kept in 150-300-gallon tanks and fed one feeder fish (fathead minnows or shiners) per gar daily. For fin regeneration studies, individuals between 20-27 cm in total length and between 263-298 dpf were anesthetized with 160mg/L MS-222 (Sigma), tagged with uniquely numbered Floy tags, and the left pectoral fin was amputated proximal to all fin rays at the endoskeleton level. Fins were fixed in 2% PFA in PBS and stored in 80% ethanol. Gars were returned to their tanks, monitored and fed daily. A total of 15 fish were sampled for a total of 15 amputations. Pectoral fin regeneration was documented with a Nikon D7100 DSLR camera and a 40mm macro lens. Fish were sampled at various stages of regrowth following euthanasia in 300mg/L MS-222 (Sigma).

### Library preparation and Illumina sequencing

For transcriptome or qPCR, total RNA extraction from different tissues was achieved using TRIzol Reagent (Life Technologies). A two step protocol, with the RNeasy Mini Kit (Qiagen) and DNaseI treatment (Qiagen), was used to purify the RNAs and remove residual DNA. The SuperScript III First-Strand Synthesis SuperMix (ThermoFisher Scientific) was used in the reverse transcription of purified total RNA (0.3 or 0.5 mg). *Polypterus* and axolotl reference transcriptomes and transcript abundance estimation were obtained from the sequencing of three blastemas, and three non-regenerating tissue libraries, performed on an Illumina 2500 Hiseq platform with 100 bp paired-end reads (PRJNA480693 and PRJNA480698).

### Bioinformatic analysis

*Polypterus* and axolotl reference transcriptomes were assembled *de novo* using Trinity with default parameters^28^ (Supplementary Fig. 3a-d). For each run, all read datasets were mapped on reference transcriptome using CLC genomic workbench with default parameters (Qiagen). For comparison between runs, expression data per transcript were summed by human homologue gene cluster using a bash script (HHGC). As previously described^29^, the HHGCs were defined by grouping transcripts with an e-value of 10^-3^ when compared by BLASTx against Human NCBI RefSeq database (11/2016). For each HHGC, the expression was calculated in transcripts per million (TPM), and the comparison was based on t-test considering two conditions (FB/LB and NRF/NRL) with three independent biological replicates.

A list of enriched GO terms was produced using the Gene Ontology Consortium web-based tool^30,31^. Differentially expressed genes with *P* values smaller than 0.05 were ranked from highest to lowest fold change values, and the corresponding ranked list of gene symbols was used for GO enrichment analysis. GO enriched categories were significant when *P* values were 0.05 or less. Reactome pathway over-representation was assessed using the Reactome web-based analysis tool, providing a gene list as input^32,33^, and then ranking results according to the over-representation score. Venn diagrams were generated using BioVenn^34^.

### Corroboration of Axolotl DGE datasets by comparison to publicly available data

Five runs were downloaded from publicly available Axolotl RNA-seq runs^16^. Three were from RNA-seq of axolotl non-regenerating upper arm tissue (SRR2885871, SRR2885875, SRR2885873) and two of a proximal blastema (SRR2885866, SRR2885865). Each run was mapped on our axolotl reference transcriptome using CLC genomic workbench with default parameters (Qiagen), and expression data in TPM was calculated by HHGC (Supplementary Table 1).

### qPCR

Each qPCR determination was performed with three biological and three technical replicates. Expression in *Polypterus* NRF (mean value of two biological replicas) was used as a reference to obtain relative expression levels in other regeneration time points. Total RNA was extracted from *Polypterus* blastema on stages of 1dpa, 5dpa and 9dpa and also NRF. For 1dpa, a pool from left and right pectoral fins from three animals was used for each biological replica. For the other regenerating stages (5dpa and 9dpa) a pool from left and right pectoral fins of one animal was used in each biological replica. Finally, for NRF, the proximal region one pectoral fin with the rays removed was used in each biological replica. Relative expression levels were assessed at 1, 5 and 9 dpa, and compared to expression levels in NRF. All experiments were performed using GoTaq Probe qPCR Master Mix (Promega)in a final volume of 10 μl. Gene-specific oligos for qPCR assays were designed using Primer Express Software Version 3.0 (ThermoFisher Scientific) and used in a final concentration of 200 nM to each primer. qPCR reactions were performed in the StepOnePlus Real-Time PCR System (Applied Biosystems) under the following cycle conditions: 2 min at 50 °C, 2 min at 95 °C followed by 40 cycles of 15 s at 95 °C, 1 min at °60 C. Relative messenger RNA expressions were calculated using the 2^-ΔΔCT^ method^35^. ΔCTs were obtained from CT normalized with *Tubb* levels in each sample. Oligos used are provided in Supplementary Table 5.

### Statistical analysis

For each transcript and HHGC, mean TPM value between NRF/NRL and FB/LB conditions was compared with a two-tailed t-test. A transcript or HHGC is classified as differentially expressed when its fold change is superior to 2 or inferior to –2 and FDR is inferior to 0.05. GO enrichment and Reactome pathway over-representation analyses were performed using the GO Consortium and Reactome web-based tools, using Fishers Exact *P* value or a statistical (hypergeometric distribution) test, respectively. qPCR analysis data were analysed using a two-tailed Welch’s corrected t-test using GraphPad Prism version 5.0 for Windows (GraphPad Software).

## Data availability

Sequence data that support the findings of this study have been deposited in GenBank with the following BioProject accession numbers: PRJNA480693 (*Polypterus* RNA-seq data) and PRJNA480698 (axolotl RNA-seq data). The authors declare that all other relevant data supporting the findings of this study are available on request.

## Acknowledgments

This work was supported by funding from CNPq Universal Program Grant 403248/2016-7 and CAPES/Alexander von Humboldt Foundation fellowship to I.S., and a postdoctoral fellowship from CNPq to A.D. M.C.D thanks research support from the NSF (IOS # 1853949). I. B. thanks support from NIH R01OD011116 for gar experiments. We thank Osage Catfisheries Inc. and the Kahrs family for their continued support of paddlefish research. We thank Allyse Ferrara and Quenton Fontenot (Nicholls State University, LA) for gar spawns, Carrie Kozel, Brett Racicot, and Solomon David for gar husbandry at Michigan State University, and John H. Postlethwait, Trevor Enright, and the University of Oregon Aquatics Facility team for support with initial gar fin regeneration trial experiments at the University of Oregon. We thank Chris Amemiya and George Mattox for insightful comments on the manuscript.

## Author contributions

S.D., M.C.D., I.B., N.F., and I.S. designed the research; A.C.D., A.N.C., A.W.T., D.B.A., J.L., M.C.D., I.B., and I.S. performed fin and limb regeneration experiments; A.C.D., D.B.A., C.M.C., J.F.S and I.S. performed and analysed qPCR data. P.N.S., constructed libraries. S.D., M.P.S., P.N.S., N.F., A.C.D., D.B.A., J.L., and I.S. analysed transcriptome data; I.S. wrote the manuscript with input from all authors. I.S. supervised this work.

## Data and materials availability

All data is available in the main text or the supplementary materials.

