## Supplementary Figures for "Deep evolutionary origin of limb and fin regeneration"

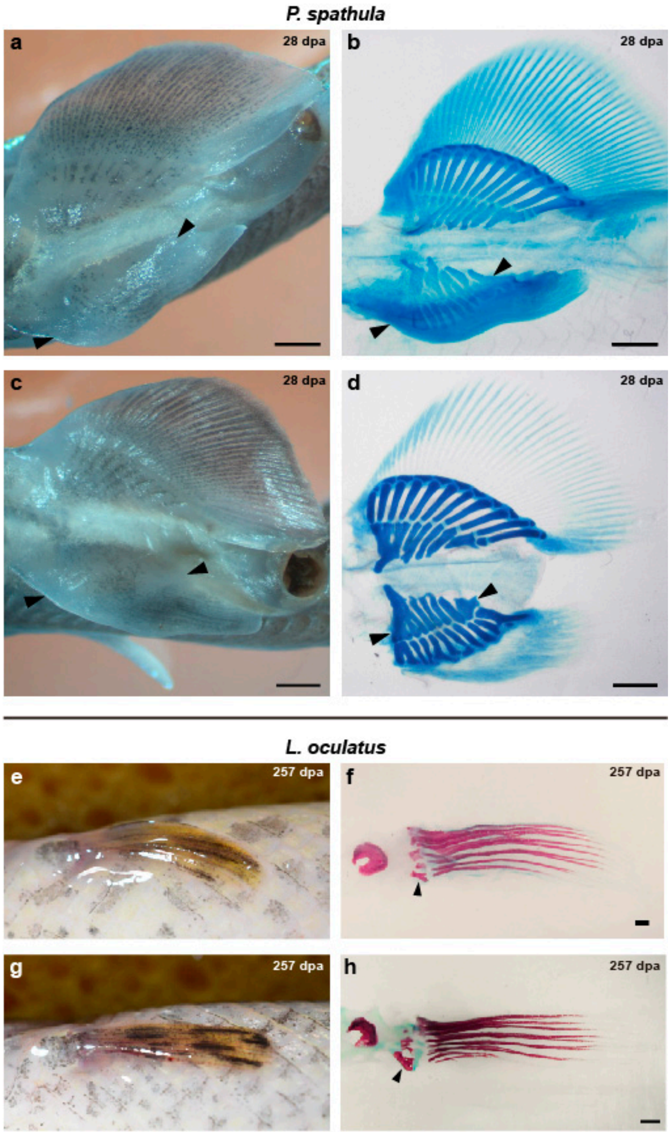


**Supplementary Fig. 1 | Additional examples of heteromorphic fin regeneration after endoskeletal amputation in *P. spathula* and *L. oculatus*.** **a** and **c**, Ventral view of specimens with regenerated right pelvic fins at 28 dpa; arrowheads denote amputation site. **b** and **d**, cleared and stained specimens showing endoskeletal regeneration distal to the amputation site. **e** and **g**, Ventral view, left pectoral fins after regeneration, at 257 dpa. **f** and **h**, Skeletal staining of regenerated fins; arrowheads denote endoskeleton. Scale bars of 1 mm. In all panels anterior is to the left.

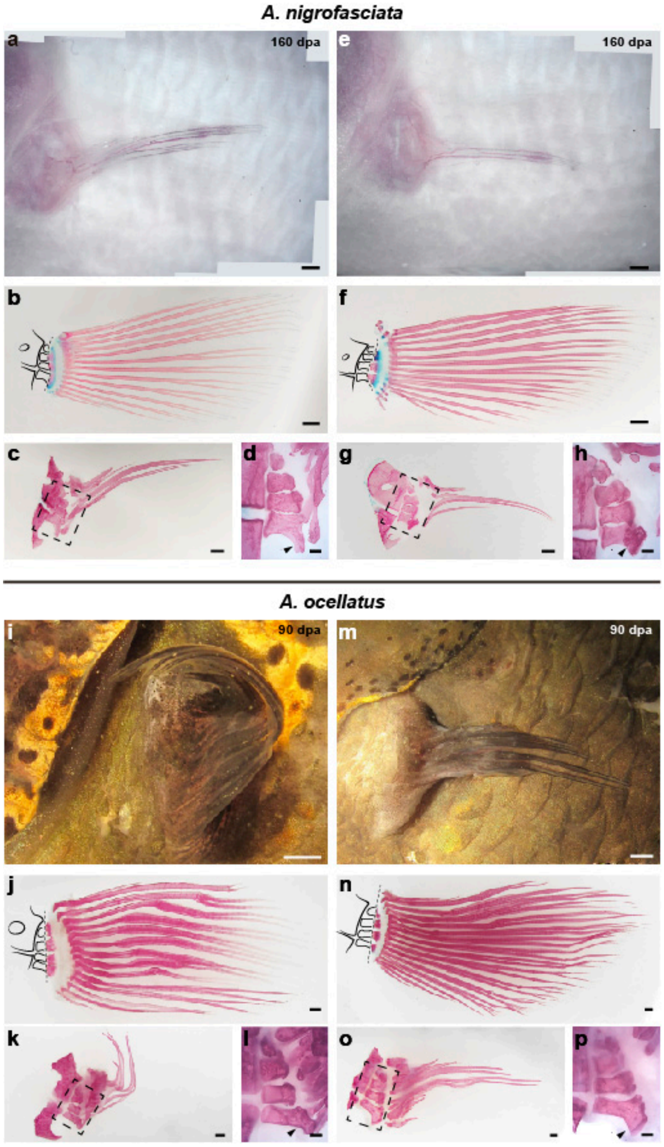


**Supplementary Fig. 2 | Additional examples of heteromorphic fin regeneration after endoskeletal amputation in cichlids. a** and **e**, White cichlid regenerated fins at 160 dpa. **b** and **f**, Skeletal staining of fins removed by amputation; dotted line denotes amputation site across the endoskeleton. **c** and **g**, Skeletal staining of regenerated fins at 160 dpa. **d** and **h**, Close-up view of regenerated endoskeleton. **i** and **m**, Oscar regenerated fins at 90 dpa. **j** and **n**, Skeletal staining of fins removed by amputation; dotted line shows amputation site across the endoskeleton. **k** and **o**, Skeletal staining of regenerated fins at 90 dpa. **l** and **p**, Close-up view of regenerated endoskeleton. Arrowheads denote endoskeleton. Scale bars of 1 mm.

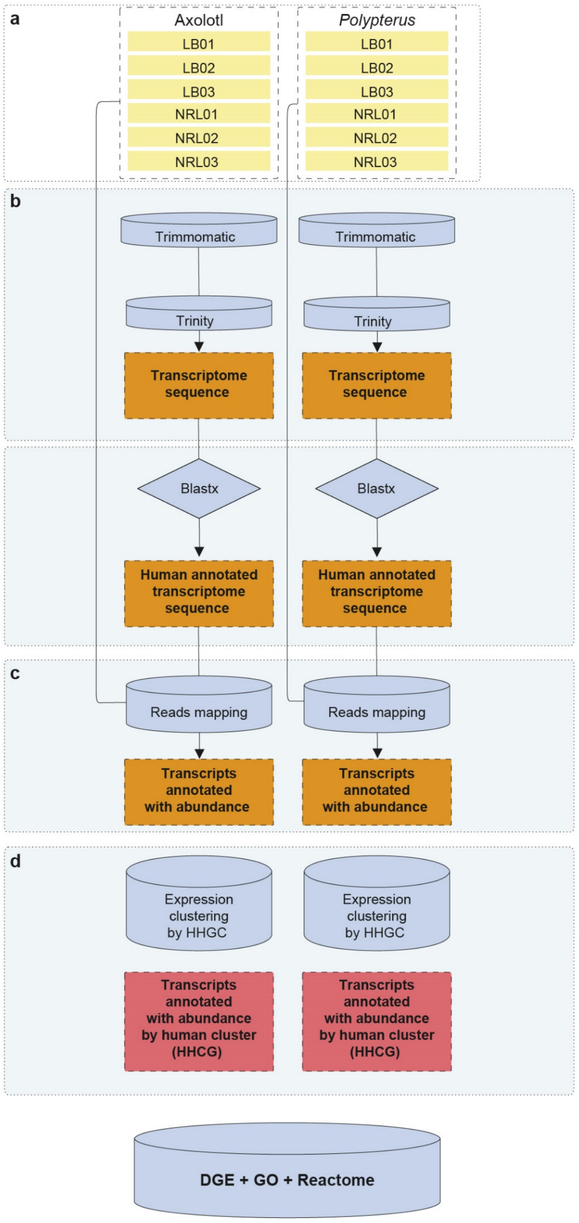


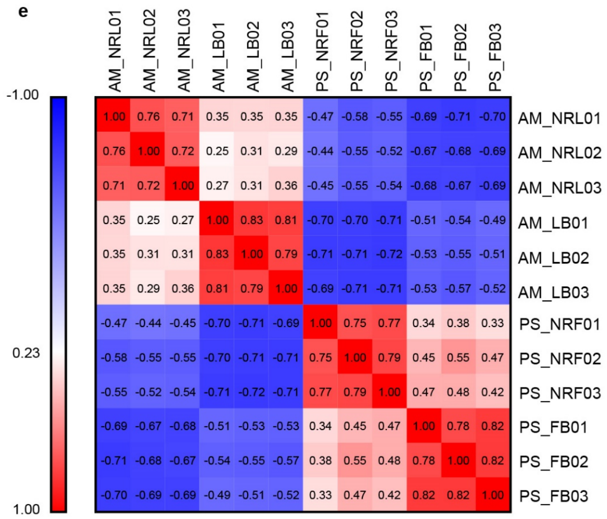


**f**

| Statistics of *de novo* transcriptome assemblies | *Polypterus* | Axolotl |
|---|---|---|
| Min contig length | 201 | 201 |
| Max contig length | 31,321 | 34,975 |
| Mean contig length | 854.38 | 895.32 |
| Standard deviation of contig length | 1292.32 | 1483.04 |
| Median contig length | 363 | 342 |
| N50 contig length | 1,933 | 2,446 |
| Number of contigs | 325,908 | 281,965 |
| Number of contigs ≥ 1kb | 66,279 | 54,567 |
| Number of contigs in N50 | 36,572 | 26,834 |
| Number of bases in all contigs | 278,449,628 | 252,448,686 |
| Number of bases in contigs ≥ 1kb | 180,689,086 | 170,336,577 |
| GC Content of contigs | 40.86 % | 45.91 % |

**Supplementary Fig. 3 | Axolotl and Polypterus RNA-seq workflow, replica correlation and reference transcriptome statistics. a-d**, Reads obtained from RNA-seq triplicates of blastema and non-regenerating tissues were used for building reference transcriptomes and subsequently mapped. Contigs were clustered by Human Homology Group Clusters (HHGC), and abundance estimated in transcripts per million for differential gene expression (DGE), gene ontology (GO) and Reactome pathway analyses. **e**, Spearman correlation coefficients among biological replicates of Axolotl (AM) non-regenerating limb (NRL), limb blastema (LB), Polypterus (PS) non-regenerating fin (NRF) and fin blastema (FB). **f**, Statistics of Axolotl and *Polypterus de novo* transcriptome assemblies.

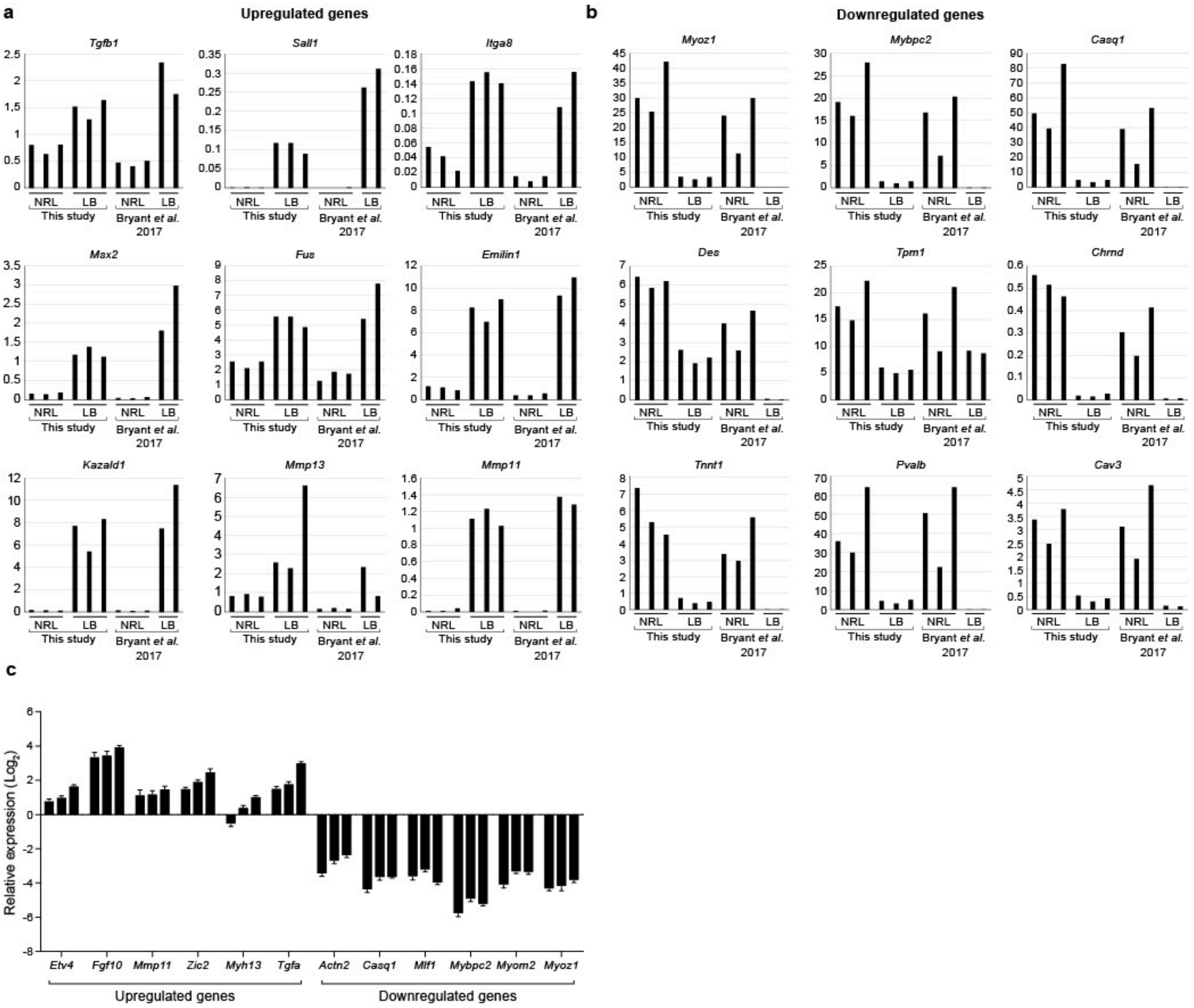


**Supplementary Fig. 4 | Corroboration of RNA-seq datasets by comparison to publicly available data and qPCR. a** and **b,** Comparison of TPM values in each run between up and downregulated genes in our study and from Bryant et al., 2017[16]. NRL, non-regenerating limb; LB, limb blastema. **c**, qPCR data of 12 genes identified as up or down regulated in *Polypterus* RNA-seq DGE dataset.

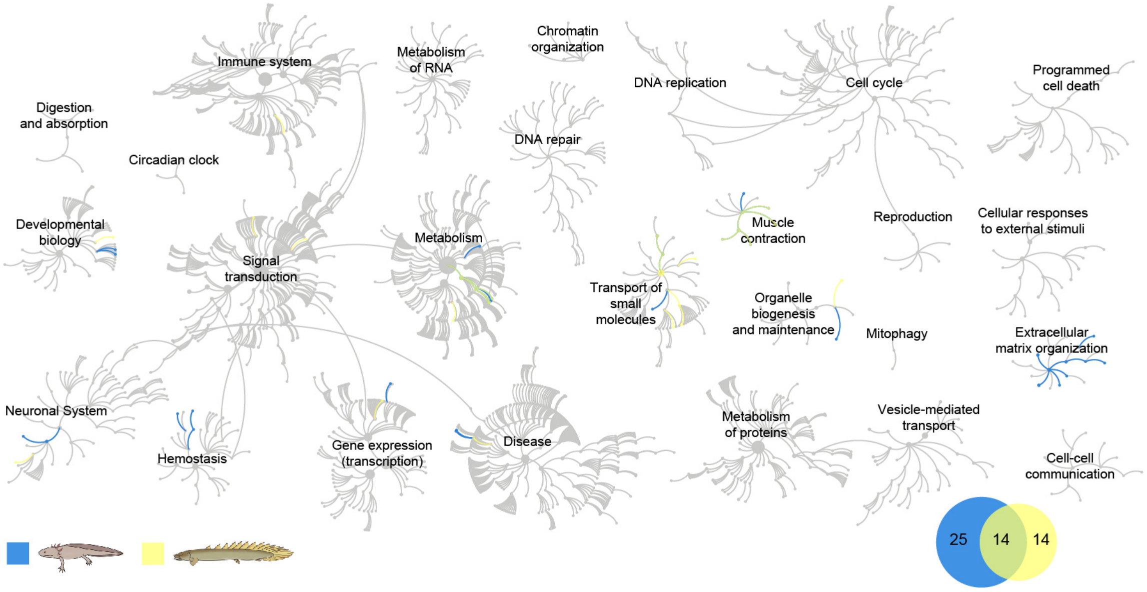


**Supplementary Fig. 5 | Over-represented pathways from downregulated gene list in axolotl and *Polypterus* blastema.** A graphical overview of Reactome pathway analysis among downregulated genes for axolotl and *Polypterus*, each of the central circles is a top-level pathway, and each step away from the center represents a lower level in the pathway hierarchy. Over-represented pathways ($P$ value<0.05) are colored in yellow (*Polypterus*), blue (axolotl) or green (overlayed pathways from both species), light grey represents pathways not significantly over-represented, an inset (bottom right) shows the Area-proportional Venn diagram of the enriched pathways in both species.

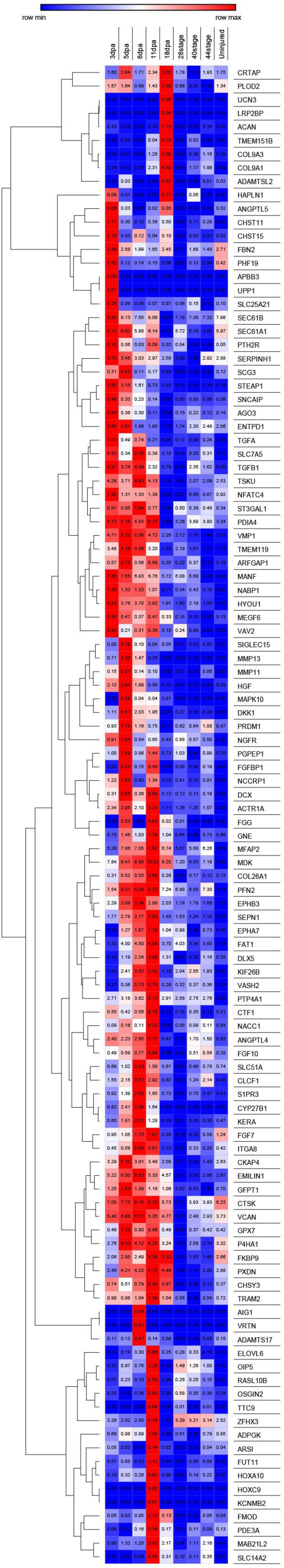


**Supplementary Fig. 6 | Genes upregulated in *Polypterus* blastema found differentially expressed during Axolotl limb regeneration.** A heatmap showing 110 genes identified as upregulated in the 9dpa *Polypterus* blastema that are differentially upregulated in Axolotl blastemas when compared to developing limb buds and to unijured limb. ScRNA-seq data from regenerating blastemas (3, 5, 8, 11 and 18dpa), developing limb buds (stages 28, 40 and 44) and uninjured limb was obtained from publically available data. For each gene, the $log_2$ TPM values were used to obtain an average expression level of all cells in a given experimental stage.
